# Ovariectomy increases paclitaxel-induced mechanical hypersensitivity and reduces anti-inflammatory CD4+ T cells in the dorsal root ganglion of female mice

**DOI:** 10.1101/2022.01.13.476262

**Authors:** Diana J. Goode, Neal E. Mecum

## Abstract

Chemotherapy is often dose limiting due to the emergence of a debilitating neuropathy. IL-10 and IL-4 are protective against peripheral neuropathy, yet the cell source is unknown. Using flow cytometry, we found that naïve females had a greater frequency of anti-inflammatory CD4+ T cells in the dorsal root ganglion (DRG) than males. In response to paclitaxel, females had reduced hypersensitivity and a greater frequency of anti-inflammatory CD4+ T cells (FoxP3, IL-10, IL-4) in the DRG than ovariectomized and male mice. These findings support a model in which estrogen promotes antiinflammatory CD4+ T cells in female DRG to suppress peripheral neuropathy.

**Highlights:**

- CD4+ T cells are present in the dorsal root ganglion of naïve and paclitaxel-treated male and female mice.
- Naïve female mice have a higher frequency of CD4+ T cells in the dorsal root ganglion compared to ovariectomized female and male mice.
- Paclitaxel induces more severe mechanical hypersensitivity in ovariectomized female and male mice compared to estrogen-competent female mice.
- Paclitaxel increases pro- and anti-inflammatory CD4+ T cells in the dorsal root ganglion of both male and female mice, but the increase in anti-inflammatory T cells is more robust in female mice.
- Ovariectomy reduces cytokine-producing CD4+ T cells in the dorsal root ganglion and prevents the PTX-induced increase in cytokine-producing CD4+ T cells in the dorsal root ganglion.

**Graphical Abstract:** Image created with Biorender.com

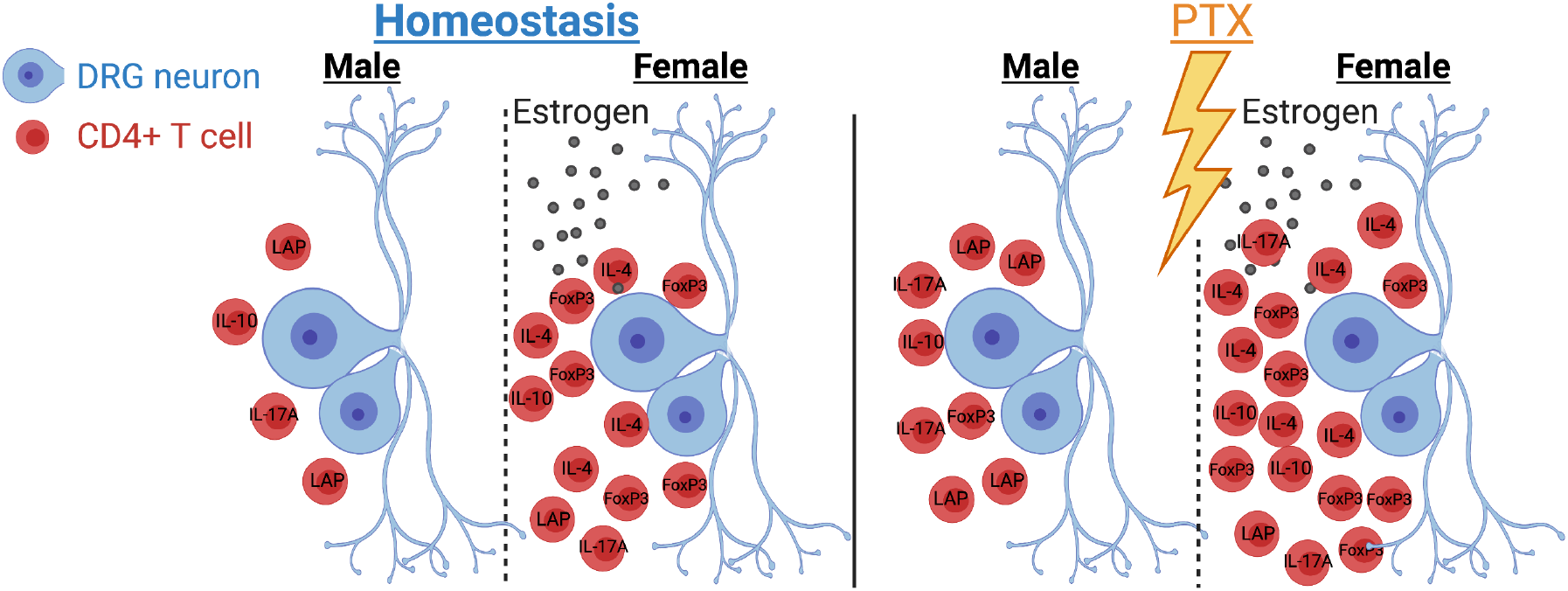

## 1. Introduction

Chemotherapeutic agents are often a life-saving cancer treatment, but the development of chemotherapy-induced peripheral neuropathy (CIPN) is a major dose-limiting toxicity resulting in intractable pain that reduces the cancer survival rate. Chemotherapeutic agents are chosen based on the type and stage of cancer. Paclitaxel (PTX) is primarily used to treat ovarian, breast, and lung cancer (Weaver, 2014); therefore, many patients are women. The prevalence of CIPN is greater than 68% (Seretny et al., 2014), with post-menopausal women (Schneider et al., 2012, Miyamoto et al., 2021) and patients taking anti-estrogens and aromatase inhibitors (Miyamoto et al., 2021) having increased risk. The onset of potentially irreversible symptoms can occur before or after completion of chemotherapy; thus, preventative measures would be invaluable.

Although the exact pathogenesis of CIPN remains elusive, one of the major off-target sites of chemotherapeutic agents is the dorsal root ganglion (DRG), which contains the sensory neurons innervating the body trunk and limbs. Due to the inefficient blood-nerve barrier, PTX accumulates in the DRG (Cavaletti et al., 2000), causing pathological changes in sensory neuron cell bodies and axons. These include neuronal hyperexcitability due to increased Nav1.7 (Li et al., 2018) and altered calcium signaling (Boehmerle et al., 2006, Li et al., 2017), and neurotoxicity due to dysfunctional axonal transport (Nakata and Yorifuji, 1999, Goshima et al., 2010) and mitochondrial damage (Flatters and Bennett, 2006). Although the central nervous system may contribute to CIPN (Omran et al., 2021), the bulk of CIPN research focuses on the peripheral nervous system because most symptoms manifest in the hands and feet (Flatters et al., 2017).

Recently, immune cells have been recognized as a major contributing factor for the resolution of CIPN (Krukowski et al., 2016, Liu et al., 2014). Although not initial thought to reside within the nervous system, both innate and adaptive resident immune cells have been found at a low abundance (0.03% - 0.5% of total DRG cells) in the DRG of naïve C57BL6/J (B6) male mice (not performed in female mice) (Liu et al., 2014). Less than a third of the immune cells in male mouse DRG (representing 0.13% of total DRG cells) are CD4+ T cells (Liu et al., 2014). Th1 and Th17 are pro-inflammatory CD4+ T cell subsets that sensitize neurons in the DRG and spinal cord to increase nociceptive-signaling (Richter et al., 2012, Vikman et al., 2003). In contrast, Forkhead box P3 (FoxP3) T regulatory cells (Tregs) and Th2 cells are anti-inflammatory CD4+ T cell subsets that suppress immune cell function through direct contact and/or anti-inflammatory cytokines. Moreover, two of these anti-inflammatory cytokines, IL-10 and IL-4, have been shown to suppress spontaneous activity from sensitized nociceptors (Krukowski et al., 2016, Laumet et al., 2020, Chen et al., 2020) and to reduce neuronal hyperexcitability (Li et al., 2018), respectively. Collectively, immune cells within the DRG orchestrate and shape immune surveillance, a process by which immune cells monitor for infection, transformation, and injury.

Recent reports demonstrate that mice lacking T cells have prolonged mechanical hypersensitivity after treatment with PTX (Krukowski et al., 2016). Liu et al. 2014 demonstrate that the frequency of CD4+ T cells (out of total leukocytes) in the DRG increases in male mice one week after PTX, at a time of peak mechanical hypersensitivity. However, the frequency of anti-inflammatory subtypes (Th2 and Treg) decreases (Liu et al., 2014). The frequency of CD4+ T cells in the DRG was not examined during the resolution of CIPN, and the effect of PTX on CD4+ T cells in female mice has never been reported. Furthermore, cytokines IL-10 (Krukowski et al., 2016) and IL-4 (Shi et al., 2018) have been shown to be protective against CIPN, yet the sources of these cytokines in the DRG after injury are unknown.

Sex-specific differences in the immune response contribute to differential susceptibilities to infectious disease, cancer, and autoimmunity (Klein and Flanagan, 2016). The main factors modulating these differences are sex hormones (Klein and Flanagan, 2016). More specifically, estrogen increases anti-inflammatory FoxP3 (Polanczyk et al., 2005), IL-10 (Yates et al., 2010), and IL-4 (Lambert et al., 2005). Moreover, females have more CD4^+^ T cells in blood and tissues (Klein and Flanagan, 2016), but it is unknown if this is true in the DRG. To have a better understanding of the function of CD4+ T cells in mouse DRG, we characterized pro- and antiinflammatory CD4+ T cell subsets in naïve and PTX-treated male, female, and ovariectomized (OVX) female mice. Our results suggest that estrogen increases anti-inflammatory CD4+ T cells in the DRG while decreasing PTX-induced mechanical hypersensitivity. Most importantly, our results propose to change current chemotherapy premedication regimens to include a component that enhances anti-inflammatory CD4+ T cells in the DRG to prevent or reduce CIPN.

## 2. Materials and Methods

### 2.1. In vivo paclitaxel administration and behavioral tests

Paclitaxel (PTX, Sigma-Aldrich-T7191) was solubilized in cremophor:ethanol 1:1 mixture and diluted 1:3 in sterile saline. A single injection of 6mg/kg PTX at a volume of 10ml/kg bodyweight was given on day 0 (as done previously (Liu et al., 2014)). von Frey and the mechanical conflict-avoidance system were used to determine mechanical hypersensitivity, and the Hargreaves Method (Hargreaves et al., 1988) was used to assess heat hyperalgesia at baseline and 3, 7, 14, 35 days post-PTX.

#### von Frey

To measure paw withdrawal frequencies to von Frey stimuli, mice were placed on a mesh screen within a plexiglass chamber. A von Frey monofilament of 3.61, which is equivalent to 0.41g force, was applied to the plantar surface of the hindpaw with enough force to bend the filament for 2-3 seconds. Paw withdrawal during and immediately after removal of the stimulus was considered a positive result. Mechanical hypersensitivity was determined by entering the response pattern in the von Frey calculator based on the up/down method (Chaplan et al., 1994).

#### Mechanical conflict-avoidance system (MCS; Coy Labs, Grass Lake, Michigan)

was adapted from previous reports (Harte et al., 2016): Mice were acclimated to the testing room and to the blinded experimenter. Mice underwent training in the MCS apparatus for 5 consecutive days. On training day 1, the alley within the MCS apparatus lacked a mechanical stimulus (no sandpaper). First, mice were placed in the dark chamber (35 lux) with a pellet of food for 5 minutes, and then placed in the light chamber with the lights off for 1 minute. After the minute expired, an aversive light (19,750 lux) was turned on and door connecting to the alley was opened. The amount of time for the whole body of the mouse to exit the light chamber and cross into the alley was recorded (latency to exit, LTE). This process was repeated 3 times for each mouse and the average value was used for the LTE. The alley of the MCS apparatus was fitted with 120 grit sandpaper for training days 2-3, and 36 grit sandpaper for training days 4-5. This same protocol using 36 grit sandpaper was used for testing days.

#### Hargreaves

Mice were placed on a glass surface (Plantar Test Analgesia Meter, Model 400 Heated Base, IITC Life Science) to measure withdrawal latency to noxious heat. A heat source (48° C) was directed towards the plantar surface of the hindpaw and turned off after hindpaw withdrawal. The elapsed time was measured to determine heat hyperalgesia. A maximum elapsed time was set to 30.1 seconds to prevent tissue damage.

### 2.2 Mice

Male, female, and OVX female (performed at 4 weeks) C57BL6/J mice were purchased from Jackson Laboratory. All mice were given food and water *ad libitum*. All experimental protocols followed National Institutes of Health guidelines and were approved by the University of New England Institutional Animal Use and Care Committee. All experiments were performed with 6–8-week-old mice. Mice were euthanized with an overdose of avertin (0.5cc of 20mg/ml) followed by transcardiac perfusion with sterile saline. This method of euthanasia is consistent with American Veterinary Medical Association (AVMA) Guidelines for the Euthanasia of Animals.

### 2.3. DRG cell isolation and flow cytometry

All DRGs (60) were collected and pooled for each mouse. DRGs were acutely dissociated as described previously (Goode and Molliver, 2021). Single cell suspensions of dissociated DRGs were passed through a 70μM cell strainer to eliminate clumps and tissue debris.

Cells were incubated at room temperature for 30 minutes with Live/Dead Fixable Violet (ThermoFisher, Waltham, MA) and 1X brefeldin A solution (BioLegend, San Diego, CA) for the intracellular cytokine staining panel. Cells were pre-incubated with anti-mouse CD16/32 (Biolegend: TruStain FcX™) for 5 minutes prior to the addition of extracellular antibodies. Cell surface antibodies (ThermoFisher: CD3ε F506, CD25 SB600, CTLA-4 FITC, PD-1 PerCP-eFluor 710, CD4 PE-Texas Red, CD44 PerCP-Cy5.5, CD28 PE-Cy7, CD62L APC, CCR7 AF700, CD69 APC-eF780) were added to the cells for 20 minutes at 4°C. Cells were washed with FACs buffer (1X PBS, 1% FBS) and fixed at room temperature for 20 minutes with Intracellular Staining Fixation Buffer (BioLegend, San Diego, CA). Cells were washed with 1X Cyto-Fast™ Perm Wash solution (BioLegend, San Diego, CA) and incubated with intracellular antibodies (Biolegend: IL-17A; ThermoFisher: IL-2 AF488, LAP PerCP-eFluor 710, Granzyme A PE, FoxP3 PerCP-Cy5.5, IL-4 PE-Cy7, Perforin APC, IL-10 AF700, IFN-γ APC-eFluor 780) for 20 minutes at room temperature. Greater than 100,000 events in the live DRG cell gate were acquired on the Beckman Coulter CytoFLEX S System B2-R3-V4-Y4 and data analyzed with FlowJo™ 10.6.1.

To ensure that our CD4+ T cell gate in the DRG was accurate, we used mouse lymph node cells as our reference sample. Lymph node (**Supplemental Figure 1A**) and DRG cells (**Supplemental Figure 1B**) were initially gated based on size (forward scatter-height, FSC-H) and granularity (side scatter-height, SSC-H) (**Supplemental Figure 1A, column 1**) to exclude cellular debris located in the bottom left corner. As the height of the cell is roughly proportional to the area, the cell height (FSC-H) was plotted against the cell area (FSC-A) (**Supplemental Figure 1A-B, column 2**) and a diagonal gate was drawn to select for single cells. As the plasma membrane of dead cells is compromised and unable to exclude the viability dye, Aqua (**Supplemental Figure 1A-B, column 3**), a gate was drawn around Aqua negative cells to eliminate dead cells from the single cell gate. CD4+ T cells were identified by using an antibody against CD3, which is expressed by all T cells, and an antibody against the co-receptor CD4, which is expressed by CD4+ T cells (**Supplemental Figure 1A-B, column 4**). For our analysis, we report cell frequencies as a percentage of CD4+ T cells, CD3+ T cells, or total DRG cells (Aqua negative cells).

### 2.4. Statistical Analysis

Group size was determined by G* Power 3 (Faul et al., 2007). All experiments were analyzed using Graphpad Prism 7 (Graphpad Software, Inc). Behavior data were checked for normality (Shapiro-Wilk), equal variances (F test), and outliers (ROUT). Behavioral data (**Figure 4, Supplemental Figure 2**) were analyzed by two-way repeated measures ANOVA with Tukey’s multiple comparisons test, and **Figure 8** behavior analyzed with a two-way ANOVA with Tukey’s multiple comparisons test. An unpaired t test was used to compare the means of CD4+ T cells in the DRG of naïve male and female mice **(Figure 1)**. A one-way ANOVA was used to compare the means of CD4+ T cells in the DRG of naïve male, female, and OVX female mice **(Supplemental Figure 3)**. Flow cytometry data (**Figure 2 - Figure 5, Supplemental Figure 4**) were analyzed by a two-way ANOVA with Tukey’s multiple comparisons test. Flow cytometry data (**Figures 6, 7**) comparing cytokine-producing CD4+ T cells at different time points within each mouse group (male, female, OVX female) were analyzed by a two-way ANOVA with Tukey’s multiple comparisons test. A two-way ANOVA with Sidak’s multiple comparisons test was used to compare individual cytokine-producing CD4+ T cell subsets over the PTX time course between male, female, and OVX female mice (**Figures 6, 7**). Results are reported as mean ± SEM with p values <0.05 to be considered significant. Degrees of freedom, F and p scores for one-way and two-way ANOVA (non-repeated measures and repeated) are presented in **Supplemental Table 1-3**.

**Figure 1.**
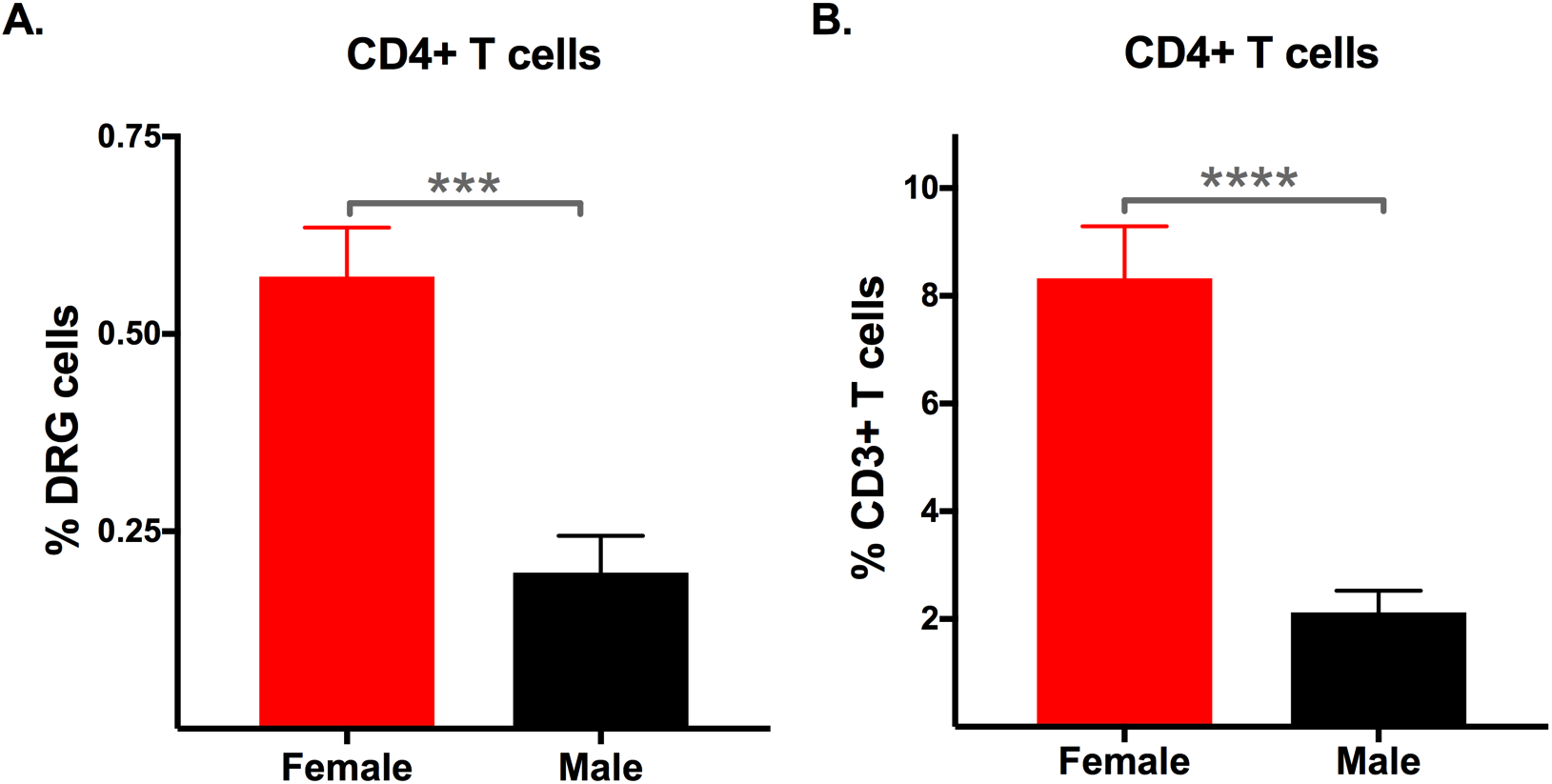
Naïve female mice have a greater frequency of CD4+ T cells in the DRG compared to naïve male mice. Frequency of **(A)** CD4+ out of total DRG cells in naïve female and male C57BL6/J (B6) mice. Statistical significance was determined by an unpaired t test (***p<0.001, n=6/sex). **(B)** Frequency of CD4+ T cells out of total CD3+ T cells in the DRG cells of naïve female and male B6 mice. Statistical significance was determined by a two-way ANOVA with Tukey’s test (****p<0.0001, n=6/sex).

**Figure 2.**
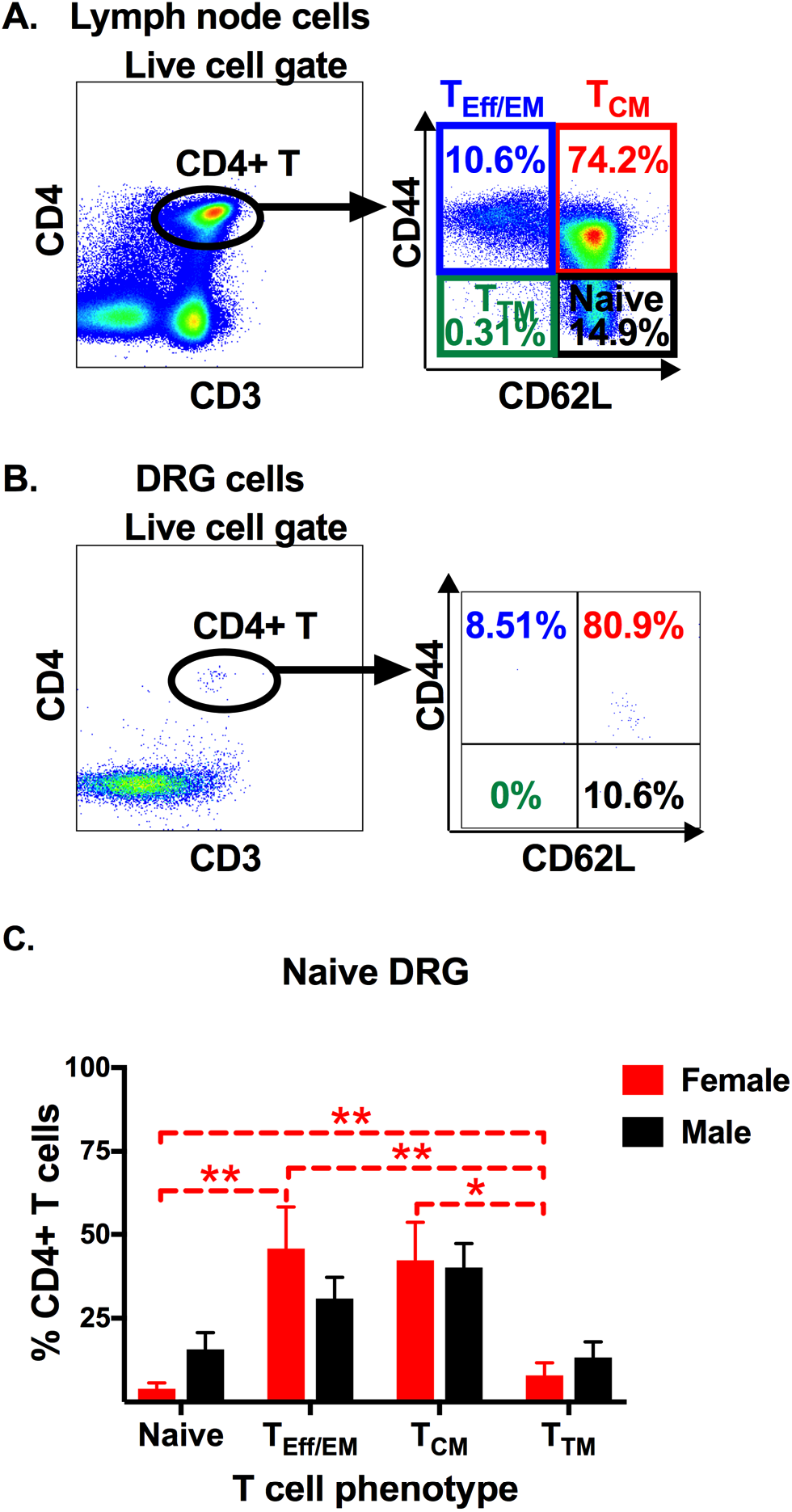
Memory and effector CD4+ T cells predominate within the DRG of female mice. The phenotype of **(A)** lymph node and **(B)** DRG CD4+ T cells was determined by plotting CD44 (memory marker) against CD62L (adhesion molecule). Based on the expression of these two markers, we quantified the frequency of naïve (CD44L-CD62L+), effector/effector memory (T_Eff/EM_: CD44+ CD62L-), central memory (T_CM_: CD44+ CD62L+), and terminally differentiated memory (T_TM_: CD44L-CD62L-) CD4+ T cells. **(C)** Frequency of naïve, T_Eff/EM_, T_CM_, and T_TM_ CD4+ T cells in the DRG of naïve female and male B6 mice (as shown in the density plot in part **B, right**). Statistical significance was determined by a two-way ANOVA with Tukey’s multiple comparisons test (*p<0.05, **p<0.01, n=6/sex).

**Figure 3.**
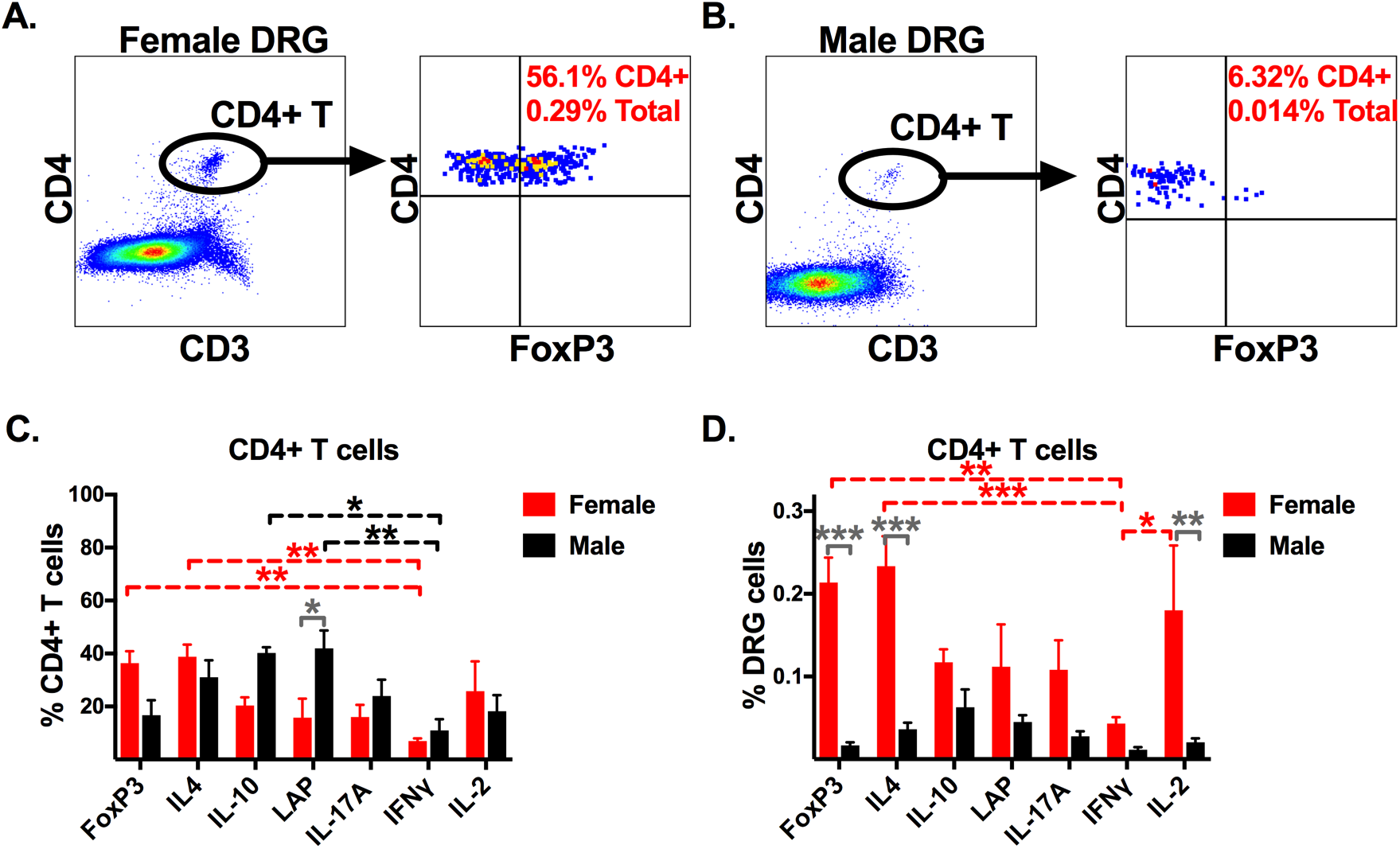
Anti-inflammatory CD4+ T cells predominate in the DRG of male and female B6 mice. Density plots of FoxP3+ CD4+ T cells from the DRG of naïve **(A)** female and **(B)** male B6 mice. Frequency of CD4+ T regulatory and cytokine-producing CD4+ T cells out of **(C)** total CD4+ T cells in the DRG and **(D)** total DRG cells. Statistical significance was determined by a two-way ANOVA with Tukey’s multiple comparisons test (*p<0.05, **p<0.01, ***p<0.001, n=6/sex). Dashed red and dashed black brackets compare cytokine-producing CD4+ T cells within females and males, respectively. Solid dark gray bracket compares CD4+ T cell subsets between males and females.

**Figure 4.**
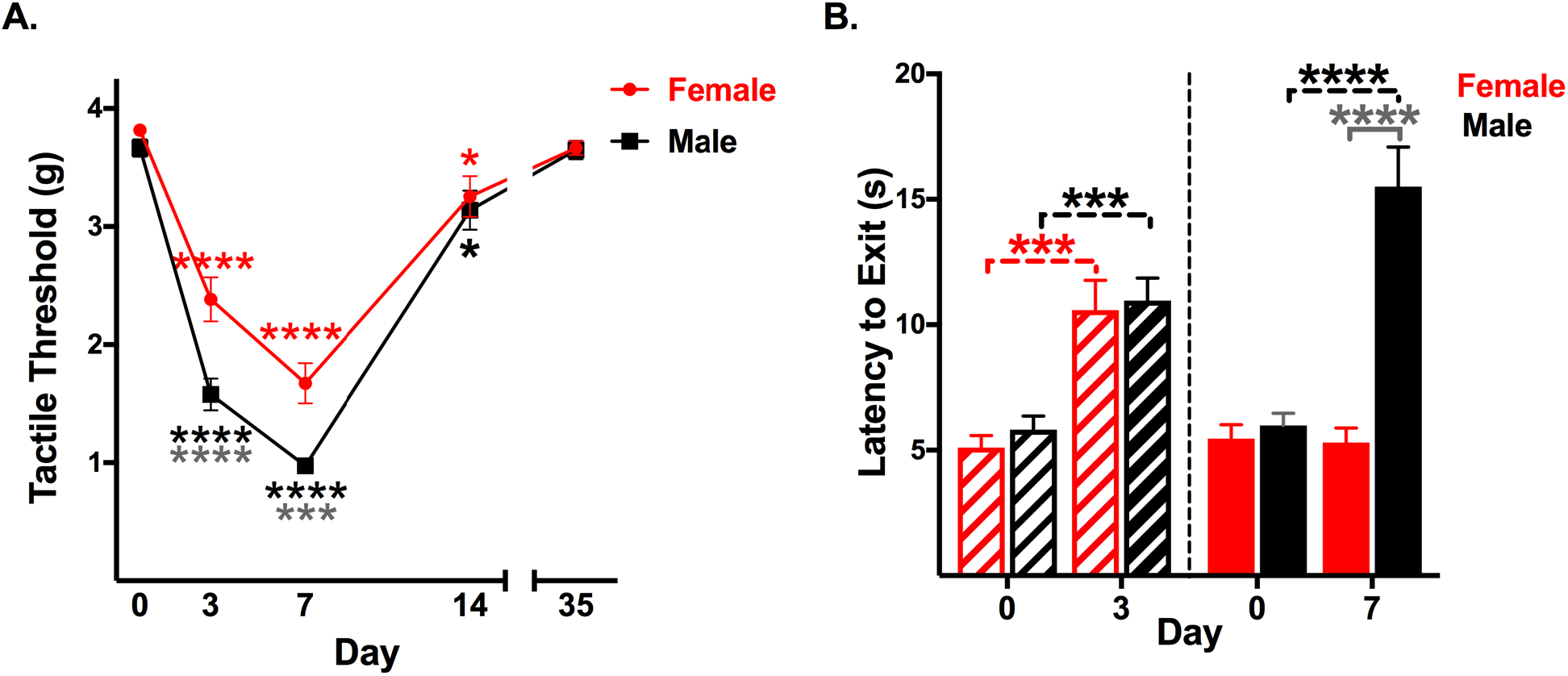
Female mice have reduced PTX-induced mechanical hypersensitivity compared to male mice. **(A)** Mechanical [tactile threshold, grams (g)] hypersensitivity assessed with von Frey filaments after 6mg/kg PTX intraperitoneal injection in female (red line) and male (black line) B6 mice. Red (female) and black (male) significance asterisks determined by a two-way repeated measures ANOVA with Tukey’s multiple comparisons test (*p<0.05, ***p<0.001, ****p<0.0001, n=27/sex) compare hypersensitivity values to baseline values for each sex. Grey significance asterisks determined by a two-way repeated measures ANOVA with Sidak’s multiple comparisons test (***p<0.001, ****p<0.0001, n=27/sex) compare hypersensitivity values between male and female mice. **(B)** Mechanical hypersensitivity assessed by mechanical conflict-avoidance system. Latency to exit [seconds (s)] the aversive lit box in PTX treated female (red) and male (black) mice 3 and 7 days post-PTX (n=13/sex; paired t test, ***p<0.001, ****p<0.0001; unpaired t test compared female mice to male mice at day 7, ****p<0.0001).

**Figure 5.**
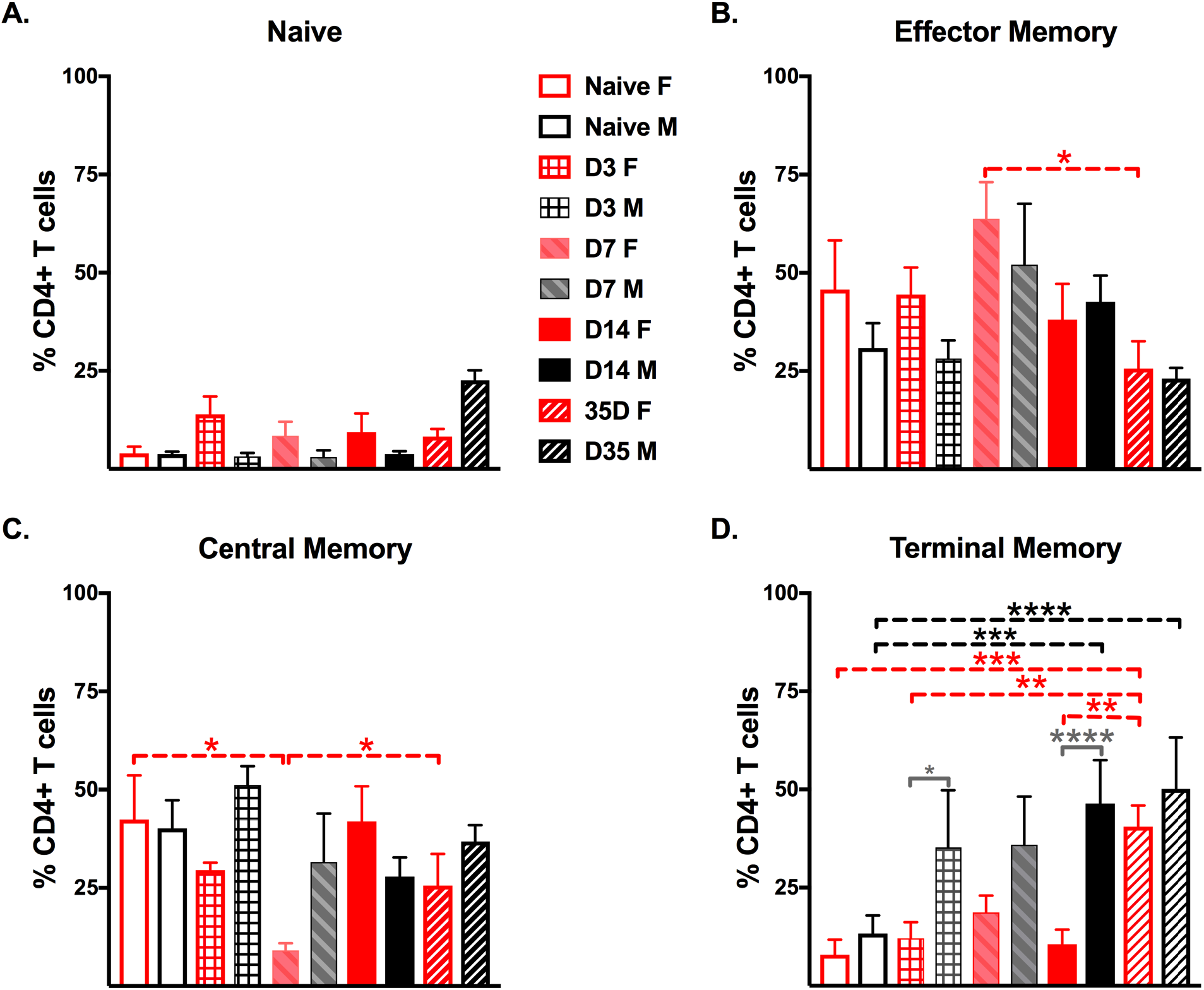
PTX increases memory CD4+ T cells in the DRG of female and male B6 mice. Frequency of **(A)** naïve (CD44L-CD62L+), **(B)** effector/effector memory (T_Eff/EM_: CD44+ CD62L-), **(C)** central memory (T_CM_: CD44+ CD62L+), and **(D)** terminally differentiated memory (T_TM_: CD44L-CD62L-) CD4+ T cells in the DRG after 6mg/kg PTX in male (black bars) and female (red bars) mice. Statistical significance was determined by a two-way ANOVA with Tukey’s multiple comparisons test (*p<0.05, **p<0.01, ***p<0.001, ****p<0.0001, n=6/sex). Dashed red and dashed black brackets compares CD4+ T cell subtypes within females and males, respectively. Solid dark gray bracket compares CD4+ T cell subtypes between males and females.

**Figure 6.**
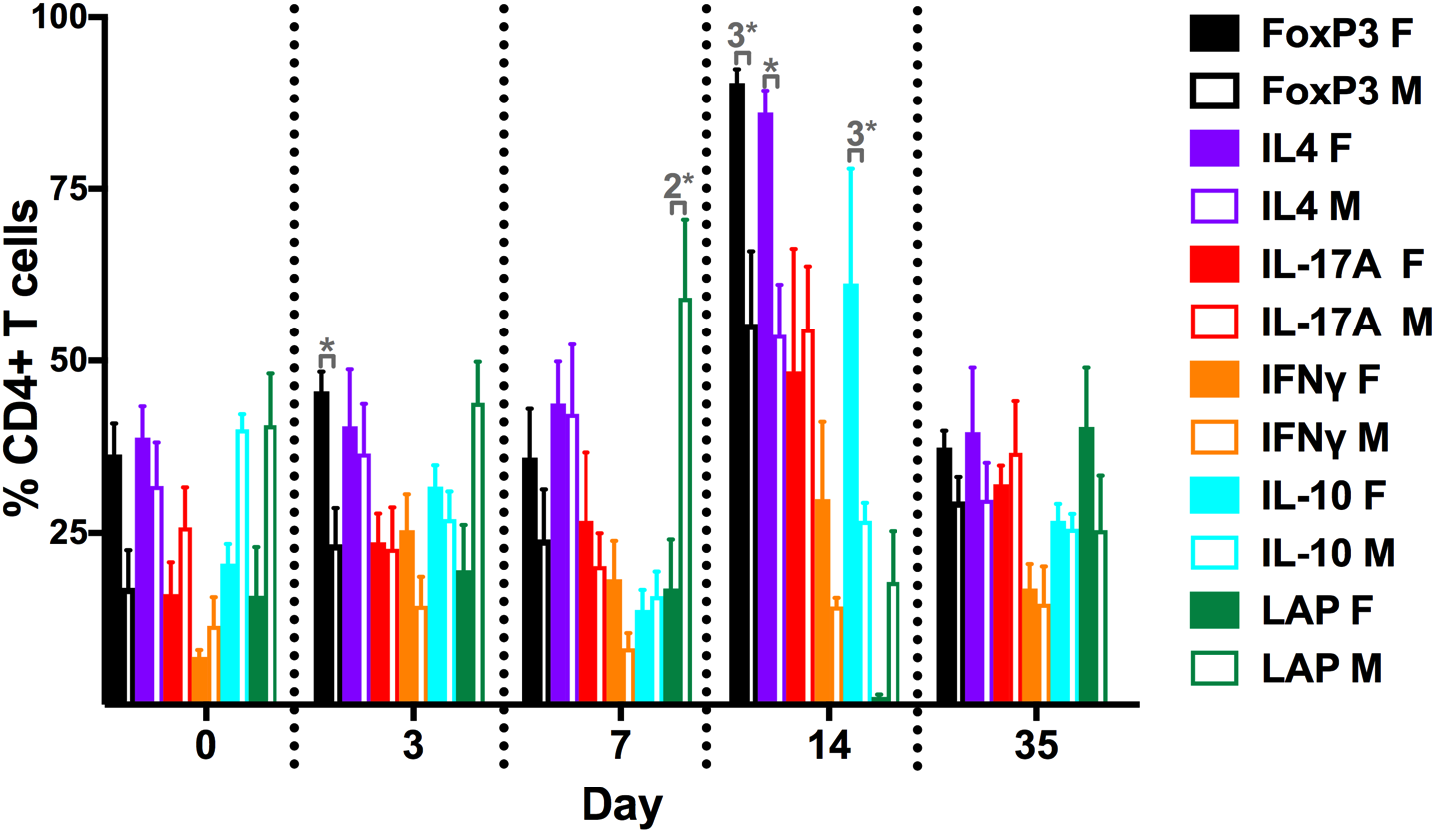
PTX robustly increases the frequency of anti-inflammatory CD4+ T cells in DRG of female mice. Frequency of CD4+ T regulatory cells and cytokine-producing CD4+ T cells in the DRG of male (open bars) and female (filled bars) mice after 6mg/kg PTX. Significance values from Tukey’s multiple comparisons test that compare different time points within each sex are listed in Table 1 (n=6/sex). Solid dark gray bracket compares cytokine-producing CD4+ T cells between males and females and significance values determined by two-way ANOVA with Sidak’s multiple comparison test (*p<0.05, 2*p<0.01, 3*p<0.001, n=6/sex).

**Figure 7.**
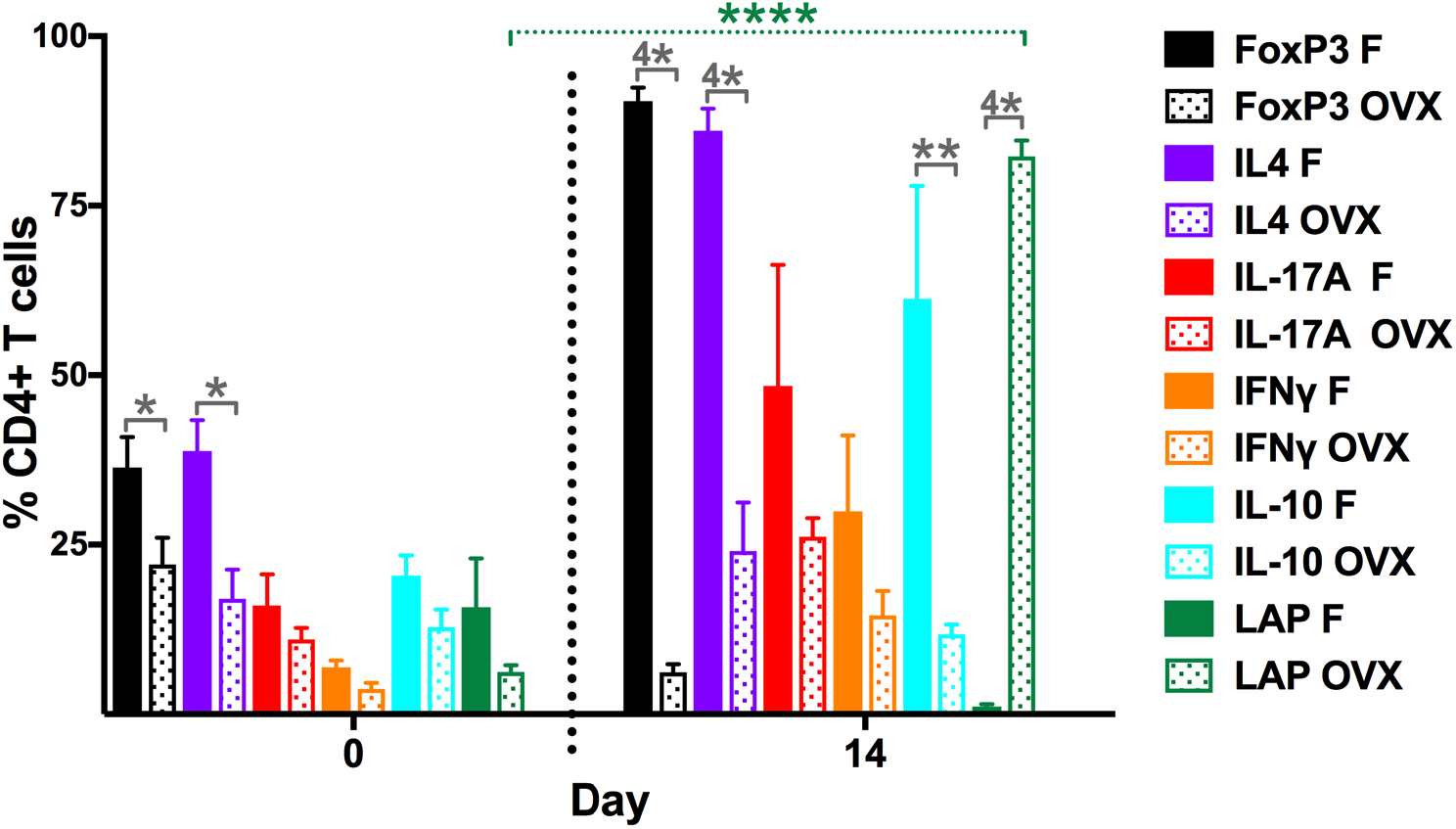
Ovariectomy reduces the frequency of anti-inflammatory CD4+ T cells in the DRG of female mice. Frequency of CD4+ T regulatory cells and cytokine-producing CD4+ T cells in the DRG of female (filled bars) and OVX (dotted bars) female mice after 6mg/kg PTX. Statistical significance determined by two-way ANOVA with Tukey’s multiple comparisons test compares CD4+ T cells in naïve OVX female mice to 14 days post-PTX (dotted green bracket, ****p<0.0001, n=5). Statistical significance determined by two-way ANOVA with Sidak’s multiple comparison test compares cytokine-producing CD4+ T cells between female and OVX female mice (*p<0.05, **p<0.01, 4*p<0.0001, n=5-6/group).

**Figure 8.**
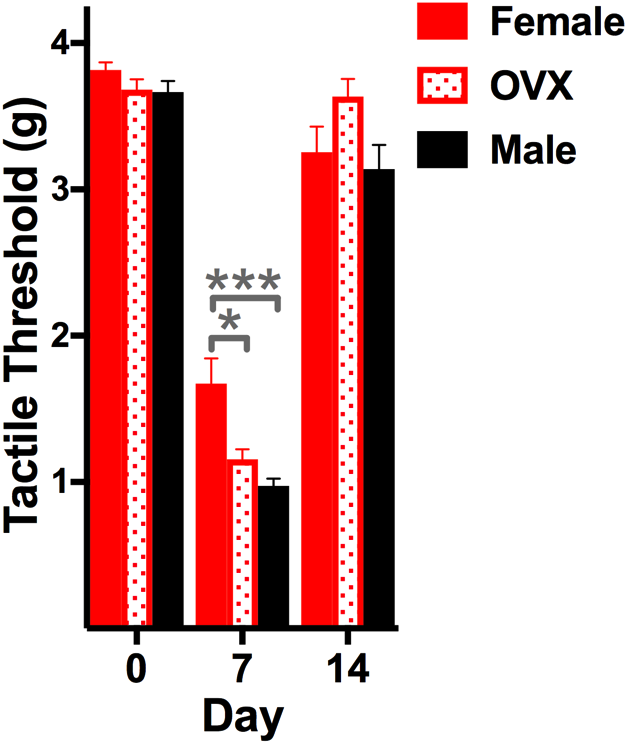
Ovariectomy increases PTX-induced mechanical hypersensitivity in female mice. Mechanical [tactile threshold, grams (g)] hypersensitivity assessed with von Frey filaments after 6mg/kg PTX intraperitoneal injection in female (solid red bar), OVX female (red dotted bar), and male (black bar) B6 mice. Gray significance asterisks determined by a two-way ANOVA with Sidak’s multiple comparisons test compares hypersensitivity values between female, OVX female, and male B6 mice (*p<0.05, 3*p<0.001, n=27 male and female; n=7-12 OVX).

## 3. Results

### 3.1 Naïve female mice have a higher frequency of CD4+ T cells in the DRG compared to male mice

Frequencies of CD4+ T cells in the blood and lymph nodes in mice have been well documented (Bio-Rad Laboratories, 2020), but frequencies of naïve/memory and cytokine-producing CD4+ T cells in the DRG of male and female mice are lacking. We found that the frequency of CD4+ T cells in mouse DRG were almost 3-fold higher in naïve female compared to naïve male (**Figure 1A,** 0.58% ± 0.06 in female versus 0.20% ± 0.11 in male, p=0.0007). Furthermore, females had a higher frequency of CD4+ T cells out of total CD3+ T cells in the DRG (8.33% ± 0.96 for female versus 2.12% ± 0.42 for male, p<0.0001) **(Figure 1B)**. The presence of CD4+ T cells in the DRG of naïve mice suggests a role for maintaining homeostasis and/or immune surveillance, a process by which immune cells monitor for infection, transformation, and injury.

### 3.2 Memory and effector CD4+ T cells predominate in the DRG of naïve female mice

The phenotype (naïve vs memory subsets) of CD4+ T cells predicts the speed and the amplitude of the cytokine response. To distinguish activated/memory CD4+ T cells from naïve CD4+ T cells, we stained with the activation marker CD44 and the adhesion molecule CD62L. We used lymph node cells **(Figure 2A)** as our reference sample to verify correct gating of naïve and memory CD4+ T cells. Consistent with the literature (Lewis et al., 2008), naïve (CD44-CD62L+) and central memory (T_CM_: CD44+ CD62L+) CD4+ T cells comprised the majority of CD4+ T cells in the lymph node. Other memory populations based on CD44 and CD62L expression include T effector/effector memory (T_Eff/EM_: CD44+CD62L-) and the terminal differentiated memory cell (T_TM_: CD44-CD62L-) (Negera et al., 2017).

We followed the same gating strategy for CD4+ T cells in the DRG **(Figure 2B)**. There were no significant differences between naïve and memory CD4+ T cells in male mouse DRG (15.68% ± 1.65 naïve, 30.87% ± 6.34 T_Eff/EM_, 40.13% ± 7.17 T_CM_, and 13.32% ± 4.62 T_TM_) **(Figure 2C)**. In contrast, T_CM_ and T_Eff/EM_ constituted the majority of CD4+ T cells in the DRG of female mice **(Figure 2C)**. CD4+ T_CM_ (42.33% ± 11.30) and T_Eff/EM_ (45.77% ± 12.47) in females were significantly greater than female naïve (3.96% ± 1.75) and T_TM_ (7.93% ± 3.82) **(Figure 2C)**. Consistent with other non-lymphoid tissues (i.e., skin, gut), the phenotype of CD4+ T cells in the DRG of female mice were effector/memory, implying a quick, robust, and local cytokine release in the DRG upon activation.

### 3.3 Anti-inflammatory CD4+ T cells predominate in the DRG of naïve male and female mice

The mere presence of CD4+ T cells in the DRG does not demonstrate functionality. Therefore, we quantified pro- and anti-inflammatory CD4+ T cells by intracellular multi-color flow cytometry. A suppressive CD4+ T regulatory (Treg) population in female (**Figure 3A**) and male (**Figure 3B**) DRG was determined by the expression of the transcription factor FoxP3. Another suppressive regulatory CD4+ T cell population was characterized by membrane-bound latency activated peptide (LAP). Naïve male mice had greater than 2.5-fold more LAP+ CD4+ T cells in the DRG than females (male 41.97% ± 6.73 compared to female 15.8% ± 7.16; p=0.0143) **(Figure 3C)**. Even though males and females had different frequencies of cytokine-producing CD4+ T cells in the DRG, the majority of CD4+ T cells in the DRG of males and females were of the antiinflammatory type. The anti-inflammatory FoxP3 (36.38% ± 4.49) and IL-4 (38.82% ± 4.54) CD4+ T cells in female DRG were present at a higher frequency than the pro-inflammatory IFN-γ (6.93% ± 1) CD4+ T cells. Likewise, the anti-inflammatory IL-10 (40.2% ± 2.18) and LAP (41.97% ± 6.73) CD4+ T cells in male DRG were present at a higher frequency than the pro-inflammatory IFN-γ (10.93% ± 4.3) CD4+ T cells. Next, we analyzed cytokine-producing CD4+ T cells out of total DRG cells. The percentages of CD4+ Tregs and IL-4 producing CD4+ T cells out of total DRG cells were 12-fold (0.213% ± 0.03 female compared to 0.017% ± 0.004 male, p=0.0002) and 6-fold (0.233% ± 0.036 female compared to 0.036% ± 0.008 male, p=0.0002, n=6) higher, respectively, in females than males **(Figure 3D)**. The T cell survival cytokine IL-2 was greater than 8-fold higher in female (0.18% ± 0.078) DRG than male (0.02% ± 0.005) **(Figure 3D)**, supporting our finding that CD4+ T cells were present in the DRG at a higher frequency than in males. Moreover, females had greater than 5-fold more anti-inflammatory Foxp3+ (0.213% ± 0.03) and IL-4+ (0.233% ± 0.036) CD4+ T cells than pro-inflammatory IFN-γ+ (0.043% ± 0.008) CD4+ T cells, which suggests that immune suppression is the default pathway in the DRG.

### 3.4 Male mice have greater PTX-induced mechanical hypersensitivity

Overall, CD4+ T cells in the DRG of naïve male and female mice were of the anti-inflammatory or CD4+ T regulatory phenotype, suggesting that the resident immune environment in the DRG is poised to prevent unnecessary immune activation to preserve tissue integrity. We administered a single intraperitoneal injection of 6mg/kg PTX to determine the extent cytokine-producing CD4+ T cells in the DRG correlated with mechanical hypersensitivity or heat hyperalgesia. von Frey testing showed that PTX induced mechanical hypersensitivity in both male and female mice on day 3 (2.38 ± 0.187 female, p<0.0001; 1.58 ± 0.136 male, p<0.0001), day 7 (1.67 ± 0.172 female, p<0.0001; 0.975 ± 0.05 male, p<0.0001), and day 14 (3.26 ± 0.174 female, p=0.0109; 3.14 ± 0.165 male, p=0.0215) post-PTX **(Figure 4A)**. Notably, female mice had reduced mechanical hypersensitivity compared to male mice day 3 (2.38 ± 0.187 female compared to 1.58 ± 0.136 male, p<0.0001) and day 7 (1.67 ± 0.172 female compared to 0.975 ± 0.05 male, p=0.0006) post-PTX **(Figure 4A)**.

Next, we used the mechanical conflict-avoidance system (MCS) (Zhou, 2014) to corroborate the PTX-induced mechanical hypersensitivity sex difference. The MCS apparatus consists of a light and dark box connected by an enclosed alley fitted with sandpaper introducing the conflicting choices of: (1) staying in the aversive brightly lit box to avoid the noxious mechanical stimulus or (2) escaping the aversive light by crossing over the noxious mechanical stimulus (sandpaper). The time to exit the light box (escape latency) was recorded. Latency to exit the brightly lit box increased 3 days following PTX treatment in male and female mice (**Figure 4B**). A sex difference was observed 7 days post-PTX, with female escape latency equivalent to baseline (pre-PTX) values (**Figure 4B, red**) whereas male mice had a delayed escape from the brightly lit box **(Figure 4B, black)**. Separate cohorts of mice were tested at 3 and 7 days post-PTX to prevent potential confounding effects of avoidance learning during the day 3 session having an impact on day 7. MCS data demonstrate that mechanical hypersensitivity is greater in males compared to females, which is consistent with results using reflexive (von Frey) measures. Of note, females recovered quicker in MCS (day 7) than von Frey (day 14), demonstrating along with other groups (Pahng and Edwards, 2018) that these tests detect overlapping but not identical measures of pain. MCS incorporates affective and motivational components of pain that reflect higher order pain processing.

Although not as severe, PTX also induced heat hyperalgesia in male and female mice 7 days post-PTX (baseline 14.88 ± 0.36 to 12.41 ± 0.65 in female, p=0.0028; baseline 15.71 ± 0.40 to 11.83 ± 0.48 in male, p<0.0001). In contrast to mechanical hypersensitivity, male and female latency response times to noxious heat were not significantly different **(Supplemental Figure 2),** suggesting that the sex-difference is unique to mechanical stimulation and not a result of a decreased pain-threshold for all nociceptive modalities.

### 3.5 PTX modulates the frequency of memory CD4+ T cells in the DRG of male and female mice

PTX did not modulate the low frequency of naïve CD4+ T cells in the DRG of male or female mice **(Figure 5A)**. However, PTX decreased the frequency of CD4+ T_Eff/EM_ in the DRG of female mice (63.75% ± 9.36 at day 7 to 25.65% ± 6.91 at day 35, p=0.0034) (**Figure 5B**). Furthermore, PTX decreased CD4+ T_CM_ cells in the DRG of female mice (naïve 42.33% ± 11.29 to 9.10% ± 1.83 at day 7, p=0.0433), but the frequency returned to baseline levels by day 14 (41.9% ± 8.95, p=0.0490) (**Figure 5C**). The frequencies of CD4+ T_TM_ in the DRG of naïve male and female mice were not significantly different. PTX increased the frequency of CD4+ T_TM_ in the DRG of both male and female mice; however, the time courses were distinct. Although there was a non-significant (p=0.0680) increase in CD4+ T_TM_ in male DRG 3 days post-PTX (naïve 13.32% ± 4.62 to 35.24% ± 14.58 at day 3), the frequency of CD4+ T_TM_ in male DRG was significantly higher than female DRG (12.11% ± 4.11, p=0.0418) (**Figure 5D**). The frequency of CD4+ T_TM_ remained elevated in the DRG of male mice on day 7 (35.89% ± 12.3), and day 14 (46.42% ±11.07, p=0.0002), which again was significantly higher than CD4+ T_TM_ in female DRG (10.57% ± 3.74, p<0.0001). The significant increase in CD4+ T_TM_ in male DRG remained high at day 35 (50.13% ± 5.39, p<0.0001) (**Figure 5D**), but was not significantly different from females as the frequency of CD4+ T_TM_ in female DRG increased 35 days post-PTX (naïve 7.93% ± 3.82 to 40.55% ± 5.39 at day 35, p=0.0003) (**Figure 5D**).

### 3.6 PTX robustly increases the frequency of anti-inflammatory CD4+ T cells in the DRG of female mice

PTX modulated the frequency of cytokine-producing CD4+ T cells in the DRG of male and female mice. Overall, PTX increased anti-inflammatory CD4+ T cells in the DRG of both male and female mice. The frequency of Foxp3+ CD4+ Tregs increased in both males (naïve 16.92% ± 5.6 to 55.35% ± 10.6, p=0.0019) and females (naïve 36.38% ± 4.49 to 90.43% ± 1.99, p<0.0001) 14 days post-PTX; however, the increase was much greater in females at both day 3 (female: 45.53% ± 2.84 to male: 23.38% ± 5.24, p=0.0409) and day 14 (female: 86.08% ± 3.26 to male: 55.35% ± 10.6, p=0.0003) post-PTX (**Figure 6**). IL-4 (naïve 31.8% ± 6.34 to 53.77% ± 7.32, p<0.0001) and IL-10 (naïve 20.42% ± 3.01 to 61.28% ± 16.65, p=0.0008) producing CD4+ T cells increased in the DRG of only female mice 14 days post-PTX, which were significantly higher than the frequencies of IL-4 (female: 90.43% ± 1.99 to male: 53.77% ± 7.32, p=0.0110) and IL-10 (female: 61.28% ± 16.65 to male: 26.78% ± 2.58, p=0.0008) in the DRG of male mice (**Figure 6**). All the frequencies that increased by day 14 in the DRG of male and female mice decreased to baseline levels 35 days post-PTX (Foxp3: male 29.58% ± 3.55 and female 37.4% ± 2.44, IL-4: female 39.63% ± 9.33, IL-10: female 26.77% ± 2.44) (**Figure 6**). Although PTX did not significantly change the frequency of LAP+ CD4+ T cells in the DRG of male and female mice, the frequency of LAP+ CD4+ T cells was much higher in male mice than female mice 7 days post-PTX (male: 59.18% ± 11.35 to female: 16.85% ± 7.21, p=0.0011). Besides increasing antiinflammatory CD4+ T cells, PTX increased the frequency of pro-inflammatory IL-17A+ CD4+ T cells in the DRG of male (naïve 25.83% ± 5.78 to 54.62% ± 9.12, p=0.0412) and female mice (naïve 16.03% ± 4.6 to 48.40% ± 17.86, p=0.0146) 14 days post-PTX (**Figure 6)**.

### 3.7 Ovariectomy reduces the frequency of anti-inflammatory CD4+ T cells in the DRG of female mice

We used OVX female mice to investigate the extent estrogen contributes to the frequency of anti-inflammatory CD4+ T cells in female DRG. Naïve OVX female mice had a lower frequency of FoxP3+ (female: 36.38% ± 4.49 to OVX: 22.09% ± 3.95, p=0.0131) and IL-4+ (female: 38.82% ± 4.54 to OVX: 17.01% ± 4.33, p=0.0109) CD4+ T cells in the DRG compared to estrogen-competent females **(Figure 7)**. In contrast to estrogen-competent mice, there was no difference between the frequencies of anti- and pro-inflammatory CD4+ T cells in the DRG of OVX female mice. To determine the extent estrogen contributes to the PTX-induced increase in anti-inflammatory CD4+ T cells in the DRG, we injected 6mg/kg PTX into OVX female mice. After 14 days, we collected the DRGs and characterized CD4+ T cells by multi-color flow cytometry. We found that ovariectomy prevented the PTX-induced increase in anti-inflammatory FoxP3+, IL-4+, and IL-10+ CD4+ T cells and pro-inflammatory IL-17A+ CD4+ T cells. The frequencies of FoxP3+ (female: 90.43% ± 1.99 to OVX: 6.16% ± 1.21, p<0.0001), IL-4+ (female: 86.08% ± 3.26 to OVX: 24.07% ± 7.17, p<0.0001), and IL-10+ (female: 61.28% ± 16.65 to OVX: 11.77% ± 1.46, p=0.0032) CD4+ T cells in the DRG of estrogen competent mice were significantly greater than the frequencies in the DRG of OVX female mice **(Figure 7)**. Besides inhibiting the PTX-induced increase of FoxP3+, IL-4+, and IL-10+ CD4+ T cells, ovariectomy drastically increased LAP+ CD4+ T cells 14 days post-PTX (naïve 6.22% ± 0.998 to 83.36% ± 1.98, p<0.0001), which was significantly greater than estrogen-competent females (female: 1.12% ± 0.356 to OVX: 82.26% ± 2.35, p<0.0001) **(Figure 7)**.

### 3.8 Ovariectomy increases PTX-induced mechanical hypersensitivity in female mice

With ovariectomy reducing anti-inflammatory CD4+ T cells in female DRG, we wanted to test whether ovariectomy increased mechanical hypersensitivity. Ovariectomy itself did not induce mechanical hypersensitivity assessed by von Frey **(Figure 8)**. However, when OVX female mice received 6mg/kg PTX, mechanical hypersensitivity increased to the level displayed by male mice. Male (0.975 ± 0.05, p=0.0001) and OVX (1.155 ± 0.07, p=0.0437) female mice had more severe mechanical hypersensitivity 7 days post-PTX compared to estrogen competent female mice (1.674 ± 0.172) **(Figure 8)**.

## 4. Discussion

### 4.1 A higher frequency of memory CD4+ T cells in the DRG of naïve female mice support an anti-inflammatory environment

With the DRG lacking a blood-nerve barrier (Jacobs et al., 1976), CD4+ T cells have been shown to infiltrate the DRG during homeostasis and injury in both humans (Esiri and Reading, 1989) and rodents (Austin et al., 2012, Liu et al., 2014, Krukowski et al., 2016). Previous flow cytometry studies did not compare the frequency of CD4+ T cells in the DRG of naive male and female mice. We found that the frequency of CD4+ T cells was almost 3-fold greater in the DRG of female mice compared to male mice **(Figure 1)**. A greater number of CD4+ T cells in females is not surprising, as many immune-related genes are on the X chromosome (Libert et al., 2010) and half the genes in activated T cells have estrogen-response elements in their promoters (Hewagama et al., 2009). Furthermore, estrogen has been documented to increase the number of CD4^+^ T cells in blood and lymphoid tissues (Klein and Flanagan, 2016). We found that the frequency of CD4+ T cells in the DRG of OVX female mice was equivalent to the frequency of CD4+ T cells in male DRG **(Supplemental Figure 3**), suggesting that estrogen drives the increased number of CD4+ T cells in female DRG. The presence of CD4+ T cells in naïve and injured DRG suggests an active process of immune surveillance, yet whether this process is antigen-dependent remains poorly understood.

Conventionally, naïve T cells produce very few cytokines, but secrete much higher amounts of cytokines upon activation by their cognate antigen in the secondary lymphoid organs (SLOs: lymph nodes and spleen). This activation process converts naïve T cells to effector T cells (T_Eff_) that have the capacity to migrate to the site of infection or injury through the loss of the adhesion molecule, CD62L. After clearance of infection or resolution of tissue injury, the majority of activated T cells die and only a few memory cells persist in peripheral tissue.

We found that the majority of CD4+ T cells in the DRG of female mice were T_Eff/EM_ and T_CM_ cells **(Figure 2)**. Although T_CM_ are thought to circulate through blood and reside in SLOs, CD4+ T_CM_ cells been shown to accumulate in the cerebrospinal fluid and choroid plexus (Kivisäkk et al., 2003) with CD62L expression providing the capacity to return to SLOs (Hengel et al., 2003), further suggesting an active role of immune surveillance in the nervous system. Interestingly, CD4+ T cells in the DRG of OVX female mice were not T_CM_ or T_Eff/EM_ but were naïve **(Supplemental Figure 4**), supporting our results that CD4+ T cells in the DRG of OVX female mice produced less cytokines **(Figure 7)**.

Similar to the gut (Izcue et al., 2006), we would expect that the default cytokine milieu within the DRG to be anti-inflammatory, as pro-inflammatory CD4+ T cell actions might result in deleterious and possibly permanent effects like nerve damage. As expected, there were more anti-inflammatory CD4+ T cells compared to pro-inflammatory T cells in the DRG of female mice **(Figure 3)**. Male DRG also had anti-inflammatory CD4+ T cells, but the frequencies were not significantly different from the frequencies of pro-inflammatory CD4+ T cells. Although both male and female DRG had anti-inflammatory CD4+ T cells, the subsets differed between the sexes. Female mice had a greater frequency of CD4+ Tregs and IL-4 producing CD4+ T cells in the DRG compared to male mice, which is consistent with the effect estrogen has on CD4+ T cells in other microenvironments — estrogen increases FoxP3 CD4+ Tregs in human blood (Singh and Bischoff, 2021) and mouse splenocytes (Polanczyk et al., 2004). Furthermore, estrogen increases IL-4 and GATA3 expression in mouse CD4+ T cells (Lambert et al., 2005). In OVX female mice, we found that ovariectomy decreased the frequency of Tregs and IL-4 producing CD4+ T cells in the DRG **(Figure 7)**, further suggesting estrogen promotes anti-inflammatory CD4^+^ T cells in female DRG under homeostatic conditions.

Male mice had a greater frequency of LAP+ CD4+ T cells in the DRG compared to female mice. LAP is a propeptide that binds to the amino terminal of TGF-β, which ensures inactivity until contact with its target cell (Nakamura et al., 2001). LAP+ CD4+ T cells mediate suppression through TGF-β (Zhong et al., 2017), and some reports suggest LAP+ T regulatory cells are more suppressive than traditional CD4+ Foxp3 Tregs (Zhong et al., 2017). LAP+ CD4+ Tregs may be a predominate mechanism for neuroimmune modulation in male mice as TGF-β receptor 1 signaling in male DRG rapidly suppresses neuronal hyperexcitability in a chronic constriction nerve injury model (Chen et al., 2015). The presence of different regulatory CD4+ T cells in male and female mice underscore the importance of sex-specific research and treatments.

### 4.2 PTX robustly increases anti-inflammatory CD4+ T cells in female DRG compared to OVX female and male DRG

Although both male and female patients develop CIPN, the immune mechanisms appear to be different, as evidenced by rodent models. Macrophages promote CIPN in both male and female mice, but TLR9 signaling in macrophages only induces pro-inflammatory cytokines (TNF-α and CXCL1) in the DRG of male mice following PTX (Luo et al., 2019). A TLR9 antagonist reverses PTX-induced mechanical hypersensitivity in male, but not female mice. Interestingly, TLR9 antagonism reverses PTX-induced mechanical hypersensitivity in T cell deficient female mouse strains (Luo et al., 2019), suggesting T cell depleted female mice revert to a male phenotype. Reversion to a male phenotype is also seen in a spared nerve injury (SNI) model of neuropathic pain (Sorge et al., 2015). Sorge et al (2015) demonstrate in a SNI model that centrally mediated mechanical hypersensitivity is dependent upon microglia in males and T cells in females; however, in the absence of T cells, females utilize glia-dependent pain pathways (Sorge et al., 2015). Recently, it has been shown that mice lacking T cells have prolonged mechanical hypersensitivity after treatment with PTX. CD8^+^ T cells could alleviate prolonged PTX-induced peripheral neuropathy in T cell deficient male and female mice (Krukowski et al., 2016, Laumet et al., 2019), but it is unclear if CD4^+^ T cell deficiency in females causes a reversion to a male phenotype. The sexual-dimorphic role of CD4^+^ T cells in the peripheral nervous system in CIPN has not been investigated. Our data demonstrate that CD4+ T cells in the DRG of estrogen competent female mice secrete anti-inflammatory cytokines (IL-10 and IL-4) **(Figure 6)** that are known to reduce CIPN.

Previous studies demonstrate that PTX increases memory T cell populations in peripheral blood of male and female patients with cancer (Wu et al., 2010, de Goeje et al., 2019). Our data shows an increase in CD4+ T_TM_ in the DRG of male and female mice, suggesting that PTX may globally enhance the frequency of memory CD4+ T cells, most likely by enhancing antigen presenting cell (APC) activation (Byrd-Leifer et al., 2001) and antigen presentation (Shurin et al., 2009). As T_CM_ cells are a renewable source of T_EM_, we found that PTX shifted the CD4+ T cell population in the DRG of female mice from T_CM_ to T_EM_ at the peak of mechanical hypersensitivity. When the mice were no longer hypersensitive, the T_EM_ /T_CM_ frequencies returned to pre-PTX levels **(Figure 6)**. While the loss of CD62L in T_EM_ is largely irreversible (Chao et al., 1997), the increase in CD4+ T_CM_ during the resolution of CIPN most likely arises from infiltration of naïve precursors, and the increase in T_TM_ is likely the result of the conversion of T_EM_ to T_TM_, as shown previously (Negera et al., 2017). These T_TM_ cells are short-lived memory cells that undergo apoptosis shortly after activation and increased proliferation (Negera et al., 2017). Therefore, the predominance of CD4+ T_TM_ cells with limited cytokine capacity in the DRG of female and male mice may contribute to increased susceptibility to peripheral neuropathy with future chemotherapeutic treatments or neuronal injury.

### 4.3 Ovariectomy increases PTX-induced mechanical hypersensitivity

Rigorous clinical studies evaluating sex as a risk factor for PTX-induced peripheral neuropathy are lacking. One report shows no sex difference (Molassiotis et al., 2019) and another reports female sex (Mizrahi et al., 2021) as a risk factor, but both were conducted in older women most likely to be post-menopausal. Studies comparing pre-versus post-menopausal women found post-menopausal women to be more susceptible to CIPN (Schneider et al., 2012, Miyamoto et al., 2021). Similarly, studies among women with gynecologic cancers found older age (hence, most likely post-menopausal) to be associated with CIPN (Bulls et al., 2019, Park et al., 2017). Furthermore, women taking anti-estrogens or aromatase inhibitors also have an increased risk of developing CIPN and symptoms tend to be more severe (Miyamoto et al., 2021). To our knowledge, no clinical study compares the magnitude or duration of PTX-induced neuropathy in pre-menopausal women to men. Our results show that PTX-induced mechanical hypersensitivity is less severe in estrogen-competent female mice than OVX female and male mice **(Figure 4)**, suggesting that estrogen is neuroprotective. Another recent report demonstrates that a sub-effective PTX dose (1mg/kg) does not induce mechanical hypersensitivity in estrogen-competent female mice but does in OVX female mice, which is prevented with daily administration of 17-β estradiol (Miyamoto et al., 2021). Our data along with others underscore the need for additional clinical studies evaluating estrogen and possibly CD4^+^ T cell deficiency as risk factors for CIPN.

## 5. Conclusions

CD4+ T cells are present in the DRG of naïve and PTX-treated male and female mice. Female mice have a greater frequency of memory/effector CD4+ T cells in the DRG than OVX female and male mice. Although the frequencies of cytokine-producing CD4+ T cells in the DRG differ between the sexes, the majority of CD4+ T cells are anti-inflammatory cells that increase after PTX administration. PTX induces a robust increase in Treg, IL-10, and IL-4 producing CD4+ T cells in the DRG of female mice, but not in OVX female or male mice. Female mice have less severe PTX-induced mechanical hypersensitivity compared to OVX female and male mice, suggesting that the estrogen-dependent anti-inflammatory CD4+ T cells in female DRG may reduce the severity of CIPN. Our work proposes that chemotherapy premedication regimens should include a component that boost anti-inflammatory CD4+ T cells in the DRG as a prophylactic measure against CIPN.

**Table 1.** CD4+ T cell results from Figure 6 two-way ANOVA with Tukey’s multiple comparison statistical analyses. The table provides Tukey’s multiple comparisons and associated significance values and p scores for two-way ANOVA analyses.

| Tukey's multiple comparisons | Significance level | P value |
| --- | --- | --- |
| **FoxP3 F** |  |  |
| 0 vs. 14 | **** | <0.0001 |
| 3 vs. 14 | *** | 0.0001 |
| 7 vs. 14 | **** | <0.0001 |
| 14 vs. 35 | **** | <0.0001 |
| **FoxP3 M** |  |  |
| 0 vs. 14 | ** | 0.0019 |
| 3 vs. 14 | * | 0.0165 |
| 7 vs. 14 | * | 0.0202 |
| **IL4 F** |  |  |
| 0 vs. 14 | **** | <0.0001 |
| 3 vs. 14 | *** | 0.0001 |
| 7 vs. 14 | *** | 0.0004 |
| 14 vs. 35 | **** | <0.0001 |
| **IL-17A F** |  |  |
| 0 vs. 14 | * | 0.0146 |
| **IL-17A M** |  |  |
| 0 vs. 14 | * | 0.0412 |
| 3 vs. 14 | * | 0.0169 |
| 7 vs. 14 | ** | 0.0072 |
| **IL-10 F** |  |  |
| 0 vs. 14 | *** | 0.0008 |
| 3 vs. 14 | * | 0.0334 |
| 7 vs. 14 | **** | <0.0001 |
| 14 vs. 35 | ** | 0.0074 |
| **LAP F** |  |  |
| 14 vs. 35 | ** | 0.0014 |
| **LAP M** |  |  |
| 7 vs. 14 | *** | 0.0006 |
| 7 vs. 35 | ** | 0.0094 |

