## Supplemental Data for "Ovariectomy increases paclitaxel-induced mechanical hypersensitivity and reduces anti-inflammatory CD4+ T cells in the dorsal root ganglion of female mice"

**Supplemental Table 1. Results from two-way ANOVA no repeated measures statistical analysis.** The table provides degrees of freedom, F and p scores for two-way ANOVA analyses.

**Supplemental Table 2. Results from two-way ANOVA repeated measures statistical analysis.** The table provides degrees of freedom, F and p scores for two-way ANOVA analyses.

**Supplemental Table 3. Results from one-way ANOVA statistical analysis.** The table provides degrees of freedom, F and p scores for one-way ANOVA analyses.

**Supplemental Figure 1. Multi-color flow cytometry gating strategy for mouse lymph node and DRG cells.** A nested gating strategy in which each gate contains the cell subset from the previous gate was used to identify CD4+ T cells. Density plots (each dot represents a single cell; red hotspots indicate a greater number of cells) of **(A)** lymph node cells and **(B)** acutely dissociated dorsal root ganglion (DRG) cells gated based on size (forward scatter-height, FSC-H) and granularity (side scatter-height, SSC-H) (column 1). Identification of single cells (from column 1 gate) was determined by plotting the cell size height against the cell size area. Single cells fall on the diagonal, as the proportion of cell height to area is roughly the same. From the single cell gate (column 2), live cells were identified based on the exclusion of the viability dye Aqua (Aqua negative). CD4+ T cells (column 4) from the live cell gate (column 3) were determined by plotting CD3 (expressed by all T cells) against CD4 (expressed by CD4+ T cells).

**Supplemental Figure 2. PTX-induces heat hyperalgesia in male and female mice.** Heat [latency, seconds (S)] hyperalgesia assessed with von Frey filaments and a radiant heat light source after 6mg/kg PTX intraperitoneal injection in female (red line) and male (black line) mice. Statistical significance between male and female sensitivity was determined by a two-way repeated measures ANOVA with Tukey's multiple comparisons test (**$p<0.01$, ***$p<0.001$, n=27/sex).

**Supplemental Figure 3. Naïve female mice have a greater frequency of CD4+ T cells in the DRG compared to naïve OVX female and male mice.** Frequency of **(A)** CD4+ out of total DRG cells in naïve female, OVX female, and male B6 mice. Statistical significance was determined by a one-way ANOVA with Tukey's multiple comparisons test (**$p<0.01$, ***$p<0.001$, n=5-6/group). **(B)** Frequency of CD4+ T cells out of total CD3+ T cells in the DRG cells of naïve female, OVX female, and male B6 mice. Statistical significance was determined by a one-way ANOVA with Tukey's multiple comparisons test (**$p<0.01$, ***$p<0.001$, n=5-6/group).

**Supplemental Figure 4. Naïve CD4+ T cells predominate within the DRG of OVX female mice.** Frequency of **(A)** naïve (CD44L- CD62L+), **(B)** effector/effector memory ($T_{Eff/EM}$: CD44+ CD62L-), **(C)** central memory ($T_{CM}$: CD44+ CD62L+), and **(D)** terminally differentiated memory ($T_{TM}$: CD44L- CD62L-) CD4+ T cells in the DRG after 6mg/kg PTX in female (red bars), OVX female (red crisscross bar), and male (black bars) mice. Statistical significance was determined by a two-way ANOVA with Tukey's multiple comparisons test (*$p<0.05$, **$p<0.01$, ****$p<0.0001$, n=5-6/group). Dashed red bracket compares CD4+ T cell subtypes within naïve OVX to 14 days post-PTX. Solid dark gray bracket compares CD4+ T cell subtypes between female, OVX female, and male B6 mice.

### Supplemental Figure 1

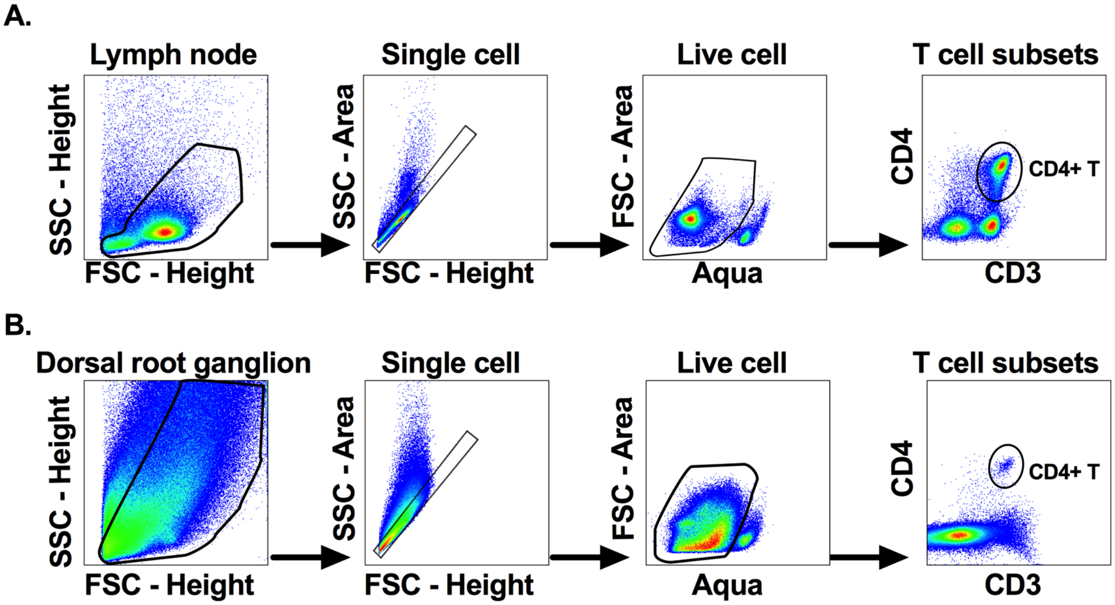

### Supplemental Figure 2

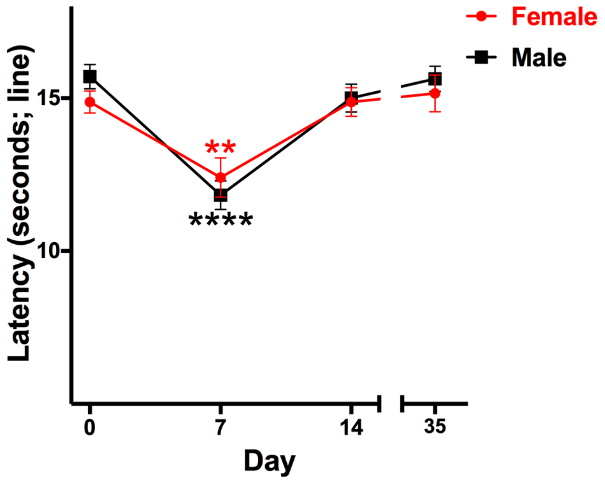

### Supplemental Figure 3

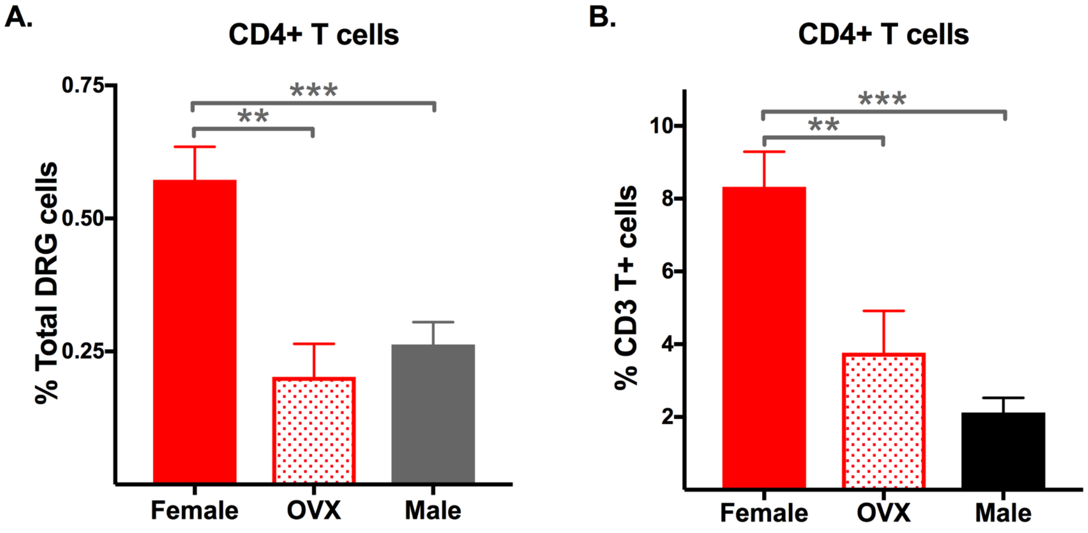

### Supplemental Figure 4

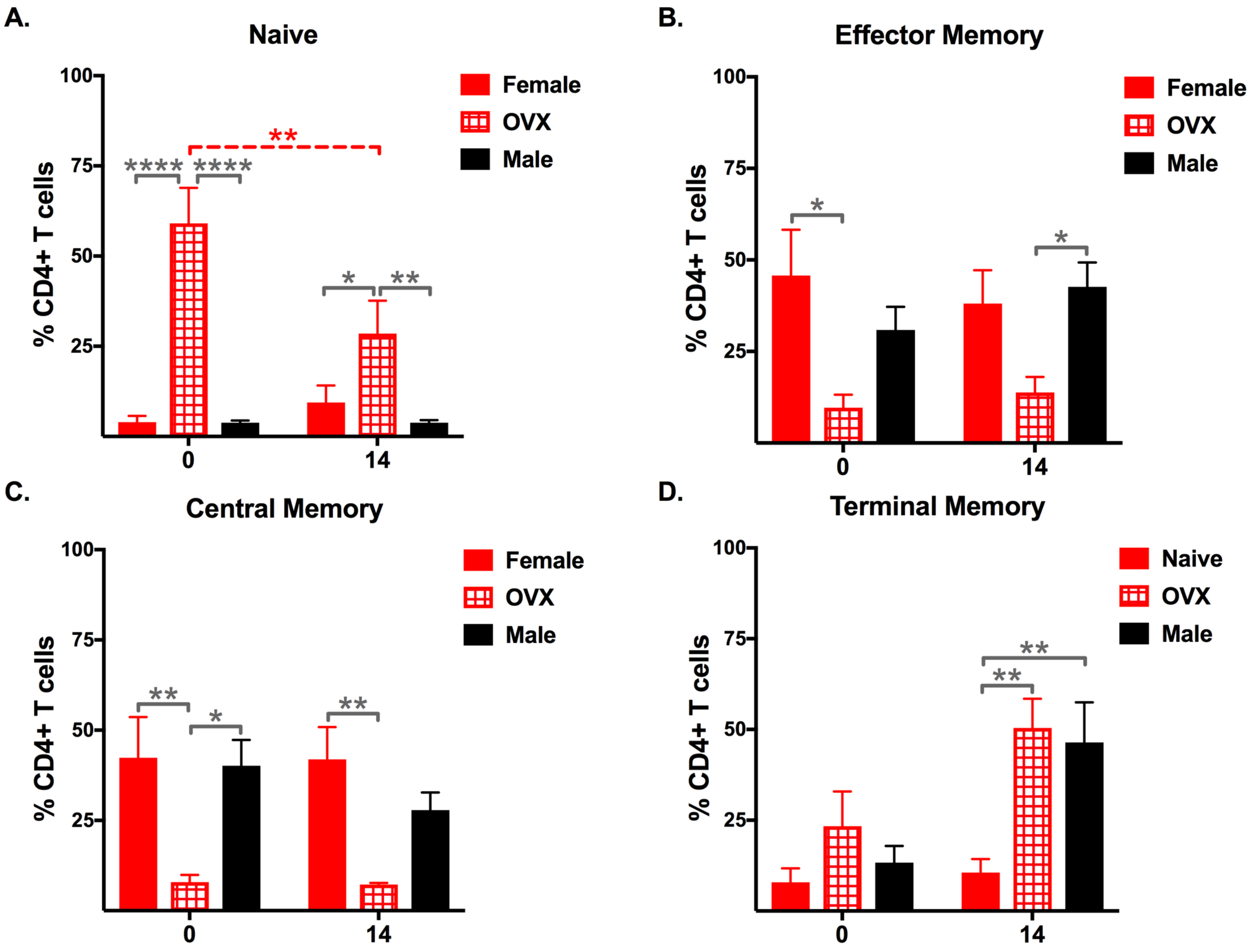

Supplemental Table 1. Results from two-way ANOVA no repeated measures statistical analysis.

| Figure # | Measurement (Units) | Cell population | | | Sex | | | Interaction | | |
|---|---|---|---|---|---|---|---|---|---|---|
| | | dfn, dfd | F | p | dfn, dfd | F | p | dfn, dfd | F | p |
| 2C | % CD4+ T cells | 3, 40 | 10.67 | <0.0001 | 1, 40 | 2.28E-07 | 0.9996 | 3, 40 | 1.196 | 0.3237 |
| 3C | % CD4+ T cells | 6, 70 | 4.338 | 0.0009 | 1, 70 | 1.114 | 0.2949 | 6, 70 | 3.962 | 0.0018 |
| 3D | % DRG cells | 6, 70 | 2.575 | 0.026 | 1, 70 | 46.71 | <0.0001 | 6, 70 | 2.617 | 0.024 |

| Figure # | Measurement (Units) | Time (0, 3, 7, 14, 35) | | | Sex/Mouse group (M, F, OVX) | | | Interaction | | |
|---|---|---|---|---|---|---|---|---|---|---|
| | | dfn, dfd | F | p | dfn, dfd | F | p | dfn, dfd | F | p |
| 5B | % CD4+ T cells | 1, 50 | 2.153 | 0.1486 | 4, 50 | 3.744 | 0.0097 | 4, 50 | 0.5089 | 0.7294 |
| 5C | % CD4+ T cells | 4, 50 | 2.627 | 0.0454 | 1, 50 | 2.817 | 0.0995 | 4, 50 | 2.286 | 0.0731 |
| 5D | % CD4+ T cells | 4, 50 | 3.995 | 0.0069 | 1, 50 | 10.74 | 0.0019 | 4, 50 | 0.9298 | 0.4543 |
| 6 | % CD4+ T cells | 4, 300 | 16.63 | <0.0001 | 11, 300 | 14.38 | <0.0001 | 44, 300 | 3.698 | <0.0001 |
| 7 | % CD4+ T cells | 1, 108 | 65.47 | <0.0001 | 11, 108 | 16.93 | <0.0001 | 11, 108 | 8.161 | <0.0001 |
| 8 | Tactile Threshold (g) | 2, 184 | 253.4 | <0.0001 | 2, 184 | 5.638 | 0.0042 | 4, 184 | 2.857 | 0.025 |
| Supp 4A | % CD4+ T cells | 1, 28 | 3.757 | 0.0627 | 2, 28 | 33.68 | <0.0001 | 2, 28 | 6.384 | 0.0052 |
| Supp 4B | % CD4+ T cells | 1, 28 | 0.171 | 0.6824 | 2, 28 | 7.53 | 0.0024 | 2, 28 | 0.7804 | 0.468 |
| Supp 4C | % CD4+ T cells | 1, 28 | 0.5465 | 0.4659 | 2, 28 | 11.35 | 0.0002 | 2, 28 | 0.4357 | 0.6511 |
| Supp 4D | % CD4+ T cells | 1, 28 | 12.52 | 0.0014 | 2, 28 | 7.909 | 0.0019 | 2, 28 | 2.608 | 0.0915 |

Supplemental Table 2. Results from two-way ANOVA repeated measures statistical analysis.

| Figure # | Measurement (Units) | Time (0, (3), 7, 14, 35) | | | Sex | | | Interaction | | |
|---|---|---|---|---|---|---|---|---|---|---|
| | | dfn, dfd | F | p | dfn, dfd | F | p | dfn, dfd | F | p |
| 4A | Tactile Threshold (g) | 4, 208 | 160.7 | <0.0001 | 1, 52 | 13.91 | 0.0005 | 4, 208 | 4.631 | 0.0013 |
| Supp 2 | Latency (seconds) | 3, 156 | 20.16 | <0.0001 | 1, 52 | 0.4212 | 0.5192 | 3, 156 | 0.7544 | 0.5214 |

Supplemental Table 3. Results from one-way ANOVA statistical analysis.

| Figure # | Measurement (Units) | Mouse group (female, male, OVX) | | |
|---|---|---|---|---|
| | | dfn, dfd | F | p |
| Supp 3A | % DRG cells | 2, 14 | 12.8 | 0.0007 |
| Supp 3B | % CD3+ T cells | 2, 14 | 14.62 | 0.0004 |
